# 3D sorghum reconstructions from depth images enable identification of quantitative trait loci regulating shoot architecture

**DOI:** 10.1101/062174

**Authors:** Ryan F. McCormick, Sandra K. Truong, John E. Mullet

## Abstract

Dissecting the genetic basis of complex traits is aided by frequent and non-destructive measurements. Advances in range imaging technologies enable the rapid acquisition of three-dimensional (3D) data from an imaged scene. A depth camera was used to acquire images of *Sorghum bicolor*, an important grain, forage, and bioenergy crop, at multiple developmental timepoints from a greenhouse-grown recombinant inbred line population. A semi-automated software pipeline was developed and used to generate segmented, 3D plant reconstructions from the images. Automated measurements made from 3D plant reconstructions identified quantitative trait loci (QTL) for standard measures of shoot architecture such as shoot height, leaf angle and leaf length, and for novel composite traits such as shoot compactness. The phenotypic variability associated with some of the QTL displayed differences in temporal prevalence; for example, alleles closely linked with the sorghum *Dwarf3* gene, an auxin transporter and pleiotropic regulator of both leaf inclination angle and shoot height, influence leaf angle prior to an effect on shoot height. Furthermore, variability in composite phenotypes that measure overall shoot architecture, such as shoot compactness, is regulated by loci underlying component phenotypes like leaf angle. As such, depth imaging is an economical and rapid method to acquire shoot architecture phenotypes in agriculturally important plants like sorghum to study the genetic basis of complex traits.

## Introduction

The rate limiting step for crop improvement and for dissecting the genetic bases of agriculturally important traits has shifted from genotyping to phenotyping, creating what is referred to as the phenotyping bottleneck (Houle et al., 2010; Furbank and Tester, 2011). Alleviating the phenotyping bottleneck for agriculturally important plants will help the world meet the increasing food and energy demands of the growing global population (Somerville et al., 2010; Alexandratos and Bruinsma, 2012; Cobb et al., 2013). Approaches to alleviate the plant phenotyping bottleneck fall into two broad categories: approaches that increase the number of individuals that can be grown and evaluated (Fahlgren et al., 2015), and approaches that predict performance *in silico* to prioritize individuals to grow and evaluate (Hammer et al., 2010; Technow et al., 2015). Both of these approaches will be instrumental for increasing the rate of crop improvement, and both approaches are facilitated by advances in image-based phenotyping; multiple plant measurements can be rapidly acquired from images, and data from image-based phenotyping approaches can also inform performance prediction (Spalding and Miller, 2013; Pound et al., 2014). As such, the development of image-based phenotyping platforms for agriculturally important plant species is a high priority for plant biology and crop improvement (Minervini et al., 2015).

The diversity of crop species and the variety of traits of interest have resulted in the development of a number of different platforms for plant phenotyping (Cobb et al., 2013; Li et al., 2014). Commercial platforms, including the Scanalyzer series from Lemnatec (http://www.lemnatec.com/products/; accessed February 2016) and the Traitmill platform from CropDesign (http://www.cropdesign.com/general.php; accessed February 2016), have gained adoption in the research community and have promoted the development of additional software (beyond that which the respective companies provide) to analyze the images produced by the platform (Reuzeau, 2007; Hartmann et al., 2011; Fahlgren et al., 2015). A variety of non-commercial platforms and methods developed by the research community also exist and have been demonstrated to perform well (White et al., 2012; Fiorani and Schurr, 2013; Sirault et al., 2013; Pound et al., 2014). Several platforms have been deployed at sufficiently large scale to examine genomic loci underlying complex traits in crop plants such as barley (Honsdorf et al., 2014), pepper (van der Heijden et al., 2012), maize (Liu et al., 2011), rice (Campbell et al., 2015), and wheat (Rasheed et al., 2014). These successful applications of image-based phenotyping to understand the genetic bases of complex crop traits represent only a small fraction of the imaging modalities and crop species available for study. Sorghum is the fifth most produced cereal crop in the world and is a promising bioenergy feedstock (Mullet et al., 2014). Recent work has demonstrated that optimization of plant canopy architecture has the potential to improve sorghum productivity (Ort et al., 2015; Truong et al., 2015). As such, we sought to develop an image-based platform to examine the genetic bases of shoot architecture traits in sorghum. While commercial products like the Scanalyzer and Traitmill systems are capable of exerting fine control and extensive automation for above-ground architecture measurements, these and other current systems did not meet our specifications for phenotyping in terms of either cost of entry, portability, output, throughput, or potential applicability in field phenotyping scenarios (Biskup et al., 2007; Sirault et al., 2013; Pound et al., 2014). Thus, we sought to develop an economical (i.e. less than 1,000 USD) image acquisition and processing pipeline capable of non-destructively assaying sorghum canopy architecture in a portable and semi-automated fashion.

Previous work has demonstrated the potential of commercial-grade depth sensors in measuring plant architecture (Chene et al., 2012; Azzari et al., 2013; Paulus et al., 2014). Therefore we used the time-of-flight depth sensor onboard a Microsoft Kinect for Windows v2 to capture depth images from multiple perspectives of individual sorghum plants, and these images were processed to construct three-dimensional (3D) representations of the imaged plants. In this manner, three replicates of 99 lines from a sorghum biparental recombinant inbred line (RIL) population were imaged at multiple timepoints during one month of development, and the images were converted to point clouds, registered, meshed, and segmented to generate 3D reconstructions of the imaged plants. Measurements from the segmented meshes and genotypes for the RIL population were used to identify quantitative trait loci (QTL) underlying shoot architecture traits. We report QTL for shoot architecture traits such as shoot height, leaf angle, and leaf length, and we demonstrate that the relative contributions to phenotypic variability of the QTL change with respect to time. We also discuss our image analysis procedures and make our code available as part of the growing body of low-cost, open-source, image-based plant phenotyping solutions.

## Materials and Methods

### Plants, greenhouse conditions, manual measurements, and image acquisition

98 recombinant inbred lines (RILs) from the BTx623 x IS3620C recombinant inbred mapping population and the two parents (Burow et al., 2011) were planted in triplicate with five seeds per pot in C600 pots of Sunshine MVP soil (BWI Companies, Inc., Texas, U.S.A.) in a College Station greenhouse on 2015-07-07. Plants were thinned to one plant per pot after germination. Plants were fertilized with Osmocote Classic (13-13-13; Everris International B.V., The Netherlands) and watered on demand. Tillers and senesced leaves were regularly removed. Each of the three replicates of the 100 RILs was grown on a separate greenhouse table, and differences in shoot morphology were visibly apparent in the population throughout development (Figures S1 and S2). Seeds for one of the RILs failed to germinate (RIL 3), leaving three replicates of 99 plants for which images were acquired. Plants were imaged at 27, 34, 39, and 44 days after planting (DAP). 15 of the plants were imaged at 62 DAP, harvested, and manually measured to compare the performance of the platform relative to standard measurement techniques. Manual measurements of leaf angle were made with a protractor, and shoot height, shoot cylinder height, leaf length, and leaf width were measured using a measuring tape. Additionally, leaf length, leaf width, and leaf area were measured using a LI-COR LI-3100C Area Meter (LI-COR, Nebraska, USA).

Image acquisition was performed using a Microsoft Kinect for Windows v2 sensor (Microsoft Corporation, Washington, USA) and the Kinect for Windows SDK (v2.0). 12 RGB and 12 depth image frames were acquired at approximately 3 second intervals and the images were saved to disk on a laptop while the Kinect for Windows v2 sensor was positioned on a tripod in front of an Arqspin 12-inch motorized turntable that rotated the imaged plant (Arqspin, Virginia, USA; Figure S3). Plants were manually transported to and from the greenhouse to the nearby imaging station. Images were transferred from the laptop to a workstation for subsequent processing.

### Processing images to acquire plant measurements

Procedures for processing images to acquire plant measurements and alternative methods that were explored are explained in File S1. Here, brief descriptions of procedures used for the reported analysis are outlined. For each plant, the point cloud contained in each depth image was automatically cleaned and registered to generate a single 3D point cloud using available open source libraries and algorithms, including OpenCV (http://opencv.org; accessed February 2016) and PCL (Fischler and Bolles, 1981; Besl and McKay, 1992; Rabbani et al., 2006; Rusu et al., 2008; Rusu and Cousins, 2011; Buch et al., 2013). This point cloud was manually inspected, acquisition and/or registration errors were manually corrected using MeshLab (Cignoni et al., 2008), and the cleaned point cloud was meshed to generate a set of polygons representing the surface of the plant using available open source software (Bernardini et al., 1999; Corsini et al., 2012; Kazhdan and Hoppe, 2013). The plant mesh was then segmented into a shoot cylinder (composed of the stem and leaf sheaths), individual leaves, and an inflorescence (when present; Figure S8). The shoot cylinder and inflorescence were manually labeled. Following this, individual leaves were segmented using an automated procedure we developed that uses supervoxel adjacency and geodesic paths across the adjacency graph to identify leaf tips and grow leaf regions (Dijkstra, 1959; Surazhsky et al., 2005; Papon et al., 2013).

Multiple measurements were automatically obtained from each mesh, both at the level of the whole plant (i.e. segmentation-independent, composite traits) and at the organ level (i.e. segmentation-dependent, organ-level traits). The traits measured are described in Table 1. Descriptions of how these traits were measured from the plant mesh are provided in File S1, and graphical depictions of selected measurements are shown in Figures S4 and S5. Additional implementation details can be found with the code base (see Code and Data Availability).

**Table 1:** Summary of the subset of traits automatically measured from the plant mesh used for the reported QTL analyses. Additional descriptions of the methods used to obtain the measurements are described in File S1.

|  | Measurement | Description of measured trait |
| --- | --- | --- |
| Composite | shoot height | vertical distance from the lowest shoot point to the highest shoot point, including leaves and the inflorescence |
|  | shoot surface area | surface area of the entire shoot |
|  | shoot center of mass | vertical distance from the lowest shoot point to the shoot's center of mass |
|  | shoot compactness | surface area of a smallest convex polyhedron that contains the entire shoot (i.e. convex hull surface area) |
| Organ-level | shoot cylinder height | vertical distance from the lowest shoot cylinder point to the highest shoot cylinder point |
|  | leaf length | length of a leaf |
|  | leaf surface area | surface area of a leaf |
|  | leaf width | width of a leaf |
|  | leaf angle | angle at which a leaf emerges from the shoot cylinder |

### QTL mapping and comparison with prior QTL studies from literature

Genotypes for the BTx623 x IS3620C RIL population were previously generated using Digital Genotyping, a restriction enzyme-based, reduced-representation sequencing assay (Morishige et al., 2013). Genotypes were called using the naïve pipeline of the RIG workflow with the GATK, and the genetic map was constructed as previously described with marker orderings relative to version 3 assembly of the sorghum reference genome, Sbi3 (DOE-JGI http://phytozome.jgi.doe.gov; accessed February 2016); this resulted in a genetic map with 10,787 markers (McKenna et al., 2010; Goodstein et al., 2012; Truong et al., 2014; McCormick et al., 2015). Both single- and multiple-QTL mapping were performed with R/qtl (Broman et al., 2003). For single-QTL mapping (i.e. testing a single-QTL model), the complete marker set of 10,787 markers was used. Measurements of a trait for each of the three replicates of a RIL were averaged; average trait values were normalized using empirical normal quantile transformation prior to QTL mapping so that the same permutation threshold would apply to all phenotype by timepoint combinations (Peng et al., 2007). A genome-wide scan under a single-QTL model for each phenotype by timepoint combination was performed (Figures S6 and S7). If any of the reported phenotype by timepoint combinations had a marker with a LOD greater than 3.28 (the 95% threshold obtained from 25,000 permutations), its LOD-2 interval (the coordinates of the flanking markers where the LODhad dropped by 2 units below the peak value) was retained. The markers’ positions (centimorgans, cM) with the largest LOD within each LOD-2 interval for each phenotype by timepoint combination were retained to initialize multiple-QTL mapping.

For multiple-QTL mapping, a subset of 1,209 markers was obtained by enforcing a minimum marker distance of 0.8 cM; significant, peak-LOD markers from single-QTL mapping intervals were added back to the set if they were dropped, resulting in 1,224 markers used for multiple-QTL mapping. The genetic coordinates of the markers with the largest LOD for each LOD-2 interval from single-QTL mapping of each phenotype by timepoint combination was used to seed model selection for multiple-QTL mapping as implemented in R/qtl (Manichaikul et al., 2009). Main effect, heavy chain, and light chain penalties (3.20, 4.38, and 1.94, respectively) for model selection were obtained as 95% thresholds from 25,000 permutations of the appropriate statistics. The multiple-QTL models with the largest penalized LOD for each phenotype by timepoint combination are reported (Tables 2, S1, and S2; Figures S6 and S7). For a given phenotype, the maximum LOD across all timepoints characterized the MLOD of the phenotype (Kwak et al., 2014). A longitudinal QTL model for each phenotype that contained QTL at the MLOD coordinates was used to generate the chromosome-wide LOD profile scans (Figures 4 and 6).

To compare QTL found in the current study with existing QTL in the literature, the physical coordinates relative to the sorghum version 1 reference assembly, Sbi1, for QTL in the BTx623 x IS3620C population were obtained; Mace and Jordan (2011) determined these physical coordinates using a consensus map and QTL identified by Hart et al. (2001) and Feltus et al. (2006). The coordinates of *Dwarf2* and *Dwarf3* were obtained from Morris et al. (2013) and Multani et al. (2003). The corresponding location of the markers in Sbi3 were obtained using Biopieces for sequence extraction and BLAST via a local instance of Sequenceserver (Hansen et al.; Altschul et al., 1997; Paterson et al., 2009; Priyam et al., 2015). Physical locations relative to Sbi3 were used as the QTL intervals for comparison with the present study.

### Code and data availability

The C++, Bash, and Python code written for image acquisition and processing, the R code written for QTL mapping, the genotype and phenotype data, and the full multiple-QTL models for each phenotype by timepoint combination can be found on GitHub at https://github.com/MulletLab/SorghumReconstructionAndPhenotyping. For each imaged plant, its depth images, a single RGB image, and the segmented mesh can be found at the DRYAD data repository (http://datadryad.org; DOI to be determined).

## Results

### 3D sorghum reconstructions from depth images

To efficiently make plant architecture measurements, a portable, economical, semi-automated image acquisition and processing pipeline was developed. Image acquisition was performed using a laptop, a tripod supporting a time-of-flight depth camera, and a turntable (Figure S3). Plants were manually transported between a greenhouse and the nearby imaging station, and, for each plant, a series of 12 depth and 12 RGB images were acquired as the plant made a 360 degree rotation on the turntable. Following acquisition, images were transferred to a workstation and processed (Figure 1).

**Figure 1:**
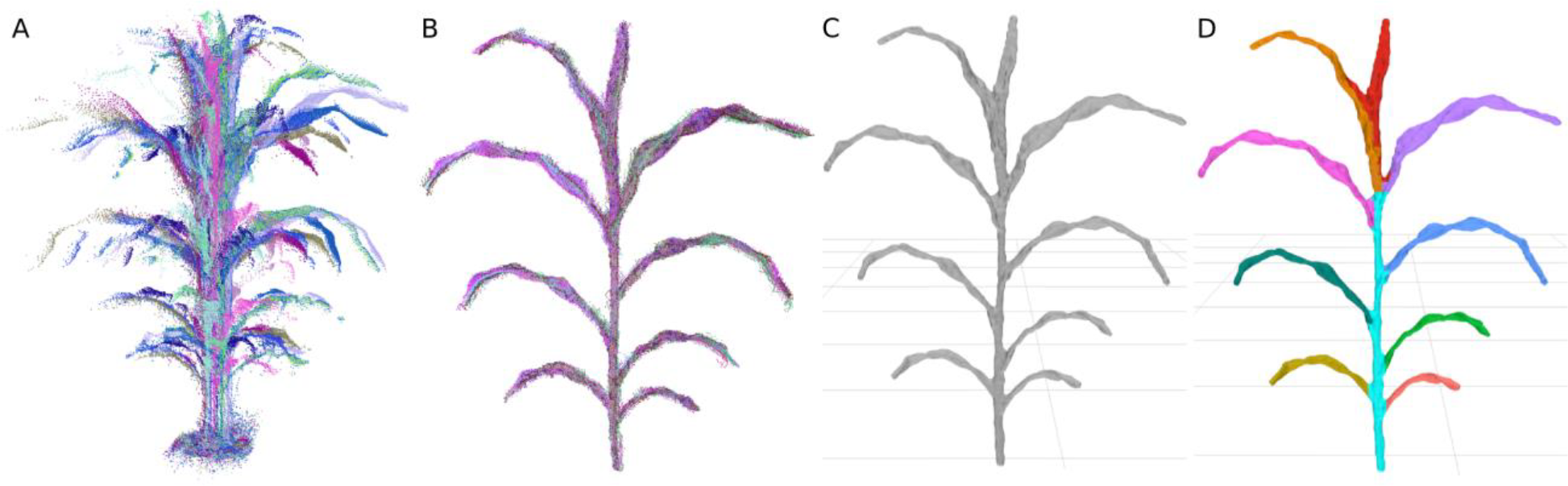
Processing of image data to segmented meshes. (A) Point clouds are sampled from multiple perspectives around the plant. (B) The point clouds are registered to the same frame and combined. (C) The combined cloud is meshed to generate a set of polygons approximating the surface of the plant. (D) The mesh is segmented into a shoot cylinder, leaves, and an inflorescence (if one exists; Figure S8), and phenotypes are automatically measured.

Most of the processing steps use generally applicable procedures available in open source libraries and software, including registration, cleaning, and meshing of the point clouds (Cignoni et al., 2008; Rusu and Cousins, 2011; Buch et al., 2013; Kazhdan and Hoppe, 2013). General solutions for segmentation of features like leaves and stems from plants, however, remain less developed, especially for 3D plant representations (Paproki et al., 2012; Paulus et al., 2013; Xia et al., 2015). Because of this, we developed a segmentation procedure for our particular application to partition the plant mesh into component parts. The final result of the semi-automated processing pipeline was a plant mesh segmented into a shoot cylinder, an infloresence (when present, Figure S8), and individual leaves with each individual leaf assigned a relative order of emergence (Figure 1).

297 plants representing triplicate plantings of 99 plants (97 RILs and the two parental lines) from the BTx623 x IS3620C sorghum mapping population were grown in a greenhouse environment (Burow et al., 2011). Because image-based phenotyping is non-destructive, the same plant can be sampled at multiple timepoints to enable change over time to be monitored. All 297 plants were imaged at four timepoints over a 17 day interval starting 27 days after planting (DAP). The four timepoints, consisting of more than 14,000 depth images and representing nearly 1,200 individual plants, were processed to segmented meshes. As such, an individual plant was represented by a timecourse of four segmented meshes, and a RIL was represented by three sets (i.e. 3 biological replicates) of an individual plant (Figure 2). A series of measurements from each mesh were then automatically acquired (Table 1 and Figure S4).

**Figure 2:**
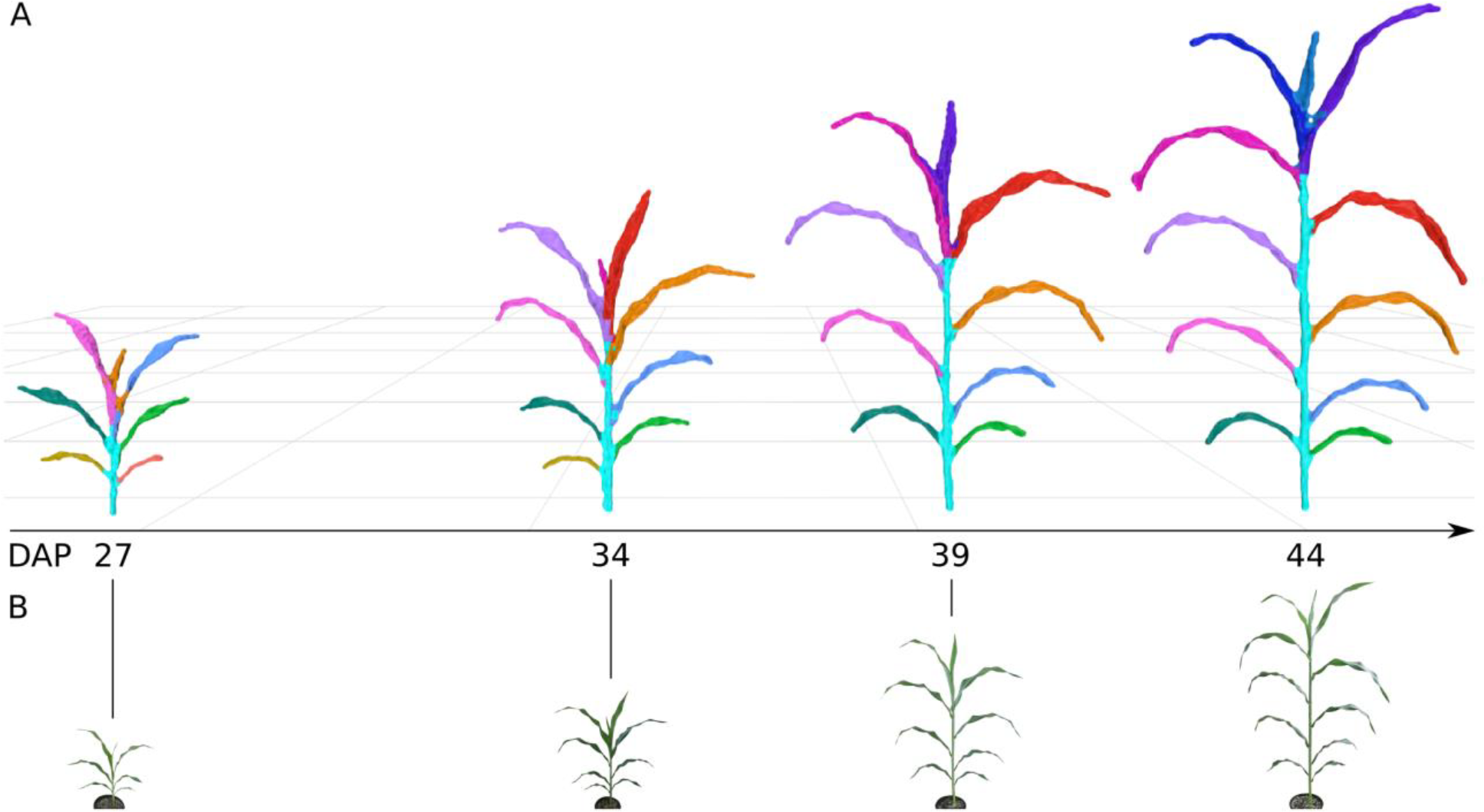
Plant growth over time. (A) Segmented meshes for replicate 3 of RIL 175 are depicted at 4 different days after planting (DAP) timepoints. Leaf colors represent individual segmented leaves; leaf colors have been manually assigned to enable tracking of the same leaf between meshes (Figure S9 depicts how color is assigned automatically by the platform). The shoot cylinder is colored cyan. Meshes are depicted at the same relative scale. (B) The corresponding RGB images that were co-acquired with the depth images; RGB images are not to scale.

To compare the measurements obtained from the image acquisition and processing platform with standard physical measurements of plant morphometric traits, 15 plants (with 140 leaves) from the experiment were imaged, and then leaf and stem measurements were obtained from harvested plants 62 days after planting. Shoot height, shoot cylinder height, leaf angle, leaf width, leaf length, and leaf area were compared. Leaf width and leaf length were measured using both a measuring tape and with a LI-COR LI-3100C Area Meter (LI-COR, Nebraska, USA), and leaf area was measured using only the LI-COR instrument. Comparisons between the measurements indicated that the image-based measurements performed at least as well as the LI-COR leaf scanning instrument for leaf width and leaf length relative to hand measurements with a measuring tape (Figure 3). The root-mean-square difference (RMSD) between manual measurements and image-based measurements for leaf length and leaf width were 7.94 cm and 1.84 cm, respectively; this indicated marginally better performance than the RMSDs between manual measurements and the LI-COR instrument for leaf length and leaf width, which were 9.41 cm and 1.94 cm, respectively. Leaf area measurements made with the depth imaging platform and with the LI-COR instrument were well correlated (Pearson product-moment correlation coefficient, ρ, of 0.92), though the image-based platform reported, on average, larger values of leaf area than the LI-COR instrument with a mean difference (MD) of 52.45 cm^2^. Leaf angle was measured with an RMSD of 9° and a ρ of 0.95 relative to hand measurements, and shoot cylinder height was measured with an RMSD of 7 cm and a ρ of 0.99. Measurements of shoot height showed the lowest correlation (ρ = 0.63 and RMSD = 11 cm) due to three outlier points; these outlier points likely represent errors in manual measurement due to the inherent difficulty in identifying the true maximum height point of the shoot in an unbiased way during manual measurement. We also note two leaf measurement outliers in both leaf length and leaf area that occurred because the image-based platform failed to fully reconstruct two of the leaves that were in the same vertical plane as the sensor. Ultimately, image-based measurements were well correlated with manual measurements and the coefficient of variation of the RMSD, CV(RMSD), for the measurements ranged from 0.07 to 0.30 (within the same range as measurements made using standard instrumentation). As such, measurements made with the phenotyping platform have utility for applications such as quantitative trait locus (QTL) mapping.

**Figure 3:**
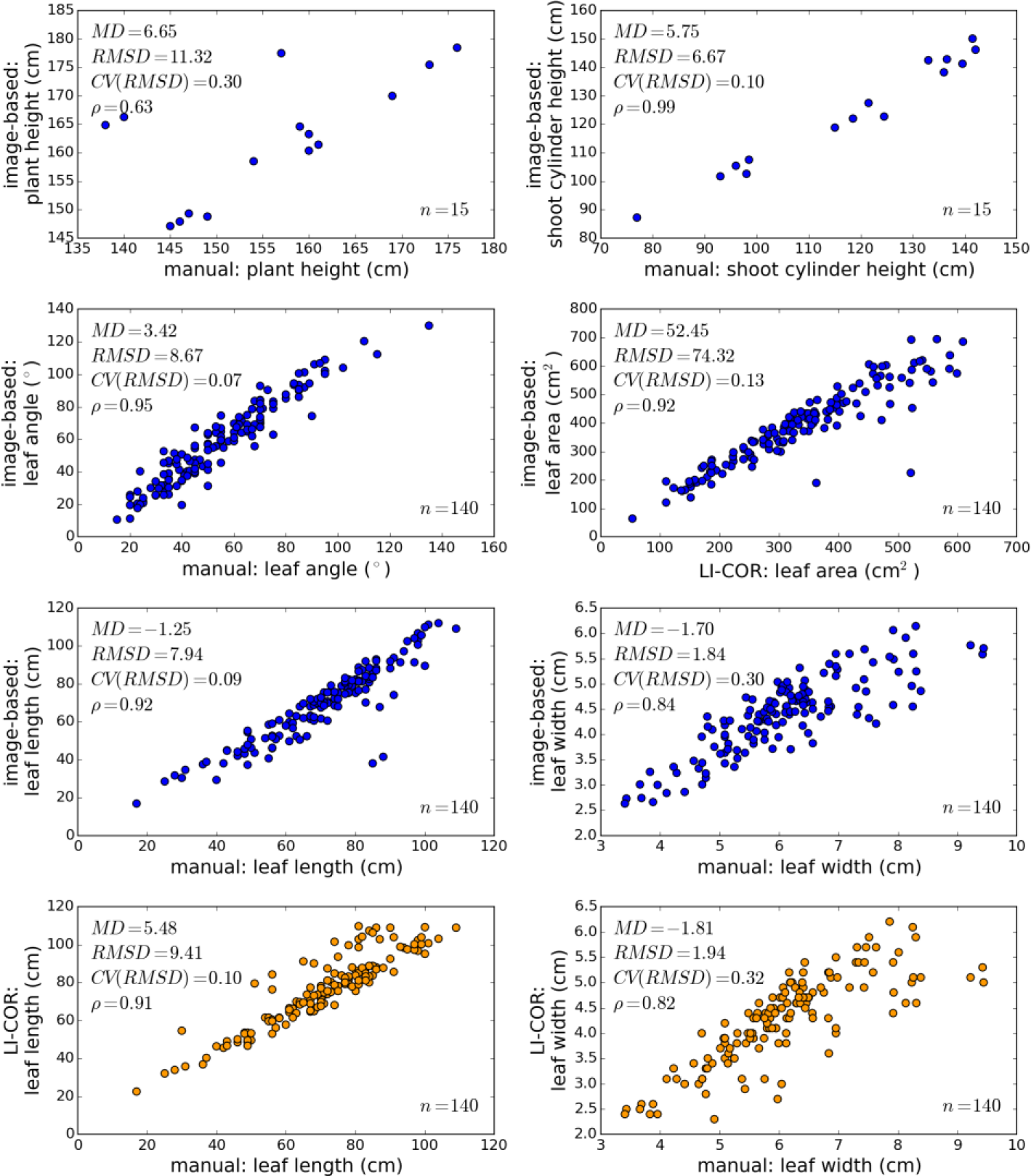
Comparison of image-based measurements with measurements made using standard methods. Axes represent measurements made via one of three methods: image-based measurements made from plant meshes, manual measurements made with a measuring tape or protractor, and measurements with a LI-COR LI-3100C Area Meter. Plots with an axis representing image-based measurements are colored blue; plots without an axis representing image-based measurements are colored orange. Leaf area measurements made with the platform include abaxial and adaxial leaf surfaces, so the image-based area measurements were divided by two for comparison with LI-COR measurements of area. *MD*: mean difference between measurements; *RMSD*: root-mean-square difference between measurements; *CV(RMSD)*: coefficient of variation of the RMSD given the range of data on the bottom axis; ρ: Pearson's product-moment correlation coefficient; *n*: number of samples from which the differences and coefficients were calculated.

### Genetic bases of imaged traits

To determine if the platform could be used to identify genetic loci regulating shoot architecture, measurements obtained from the plant meshes were associated with genetic data from the RIL population.Genotypes for members of the BTx623 x IS3620C RIL population were previously obtained and available to construct a genetic map for mapping quantitative trait loci (QTL) for the image-based phenotypes across multiple developmental timepoints (Morishige et al., 2013; Truong et al., 2014; McCormick et al., 2015). Measurements obtained from plant meshes were grouped into two categories: organ-level measurements and composite measurements. Organ-level measurements are segmentation-dependent and measure organ-level plant architecture, such as leaf length and shoot cylinder height; composite measurements are segmentation-independent and measure overall shoot architecture such as shoot height and shoot compactness (Table 1, Figure S3 and Figure S4).

QTL mapping of organ-level traits identified seven unique genomic intervals with significant contributions to phenotypic variability (Figure 3, Figure S5, and Table S1). A genome-wide scan under a single-QTL model was used to examine the following phenotypes across the four timepoints: the average value of leaves 3, 4, and 5 for leaf length, width, surface area, and inclination angle, and shoot cylinder height. Significant QTL identified from a genome-wide scan under a single-QTL model were used as an initial model for stepwise model traversal to identify the most likely penalized multiple-QTL model (Manichaikul et al., 2009); the overlapping LOD-2 intervals of these multiple-QTL models define unique intervals on chromosomes 3, 4, 6, 7, and 10 (Table S1).

**Figure 4:**
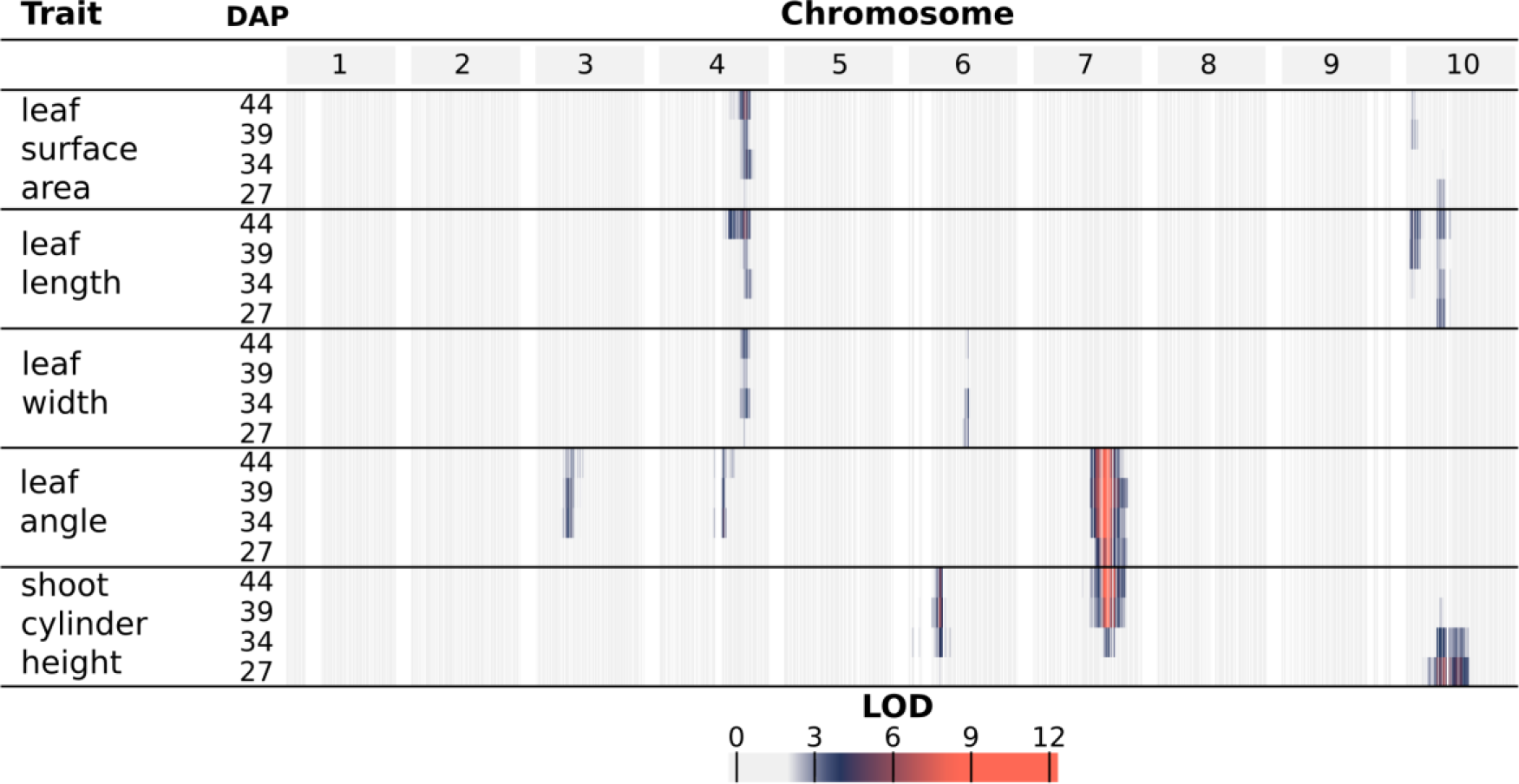
LOD profiles for organ-level traits. For each phenotype, LOD profiles are based on chromosome-wide scans of chromosomes with QTL based on the most likely multiple-QTL models found by model selection (Figure S6). Each row represents a different trait, and within each trait are four nested rows that each represents a different timepoint (days after planting; DAP). Each group of columns represents a chromosome, and each column represents a marker at its genetic position. Cells are colored by marker LOD for the phenotype at the particular timepoint.

A major source of variation in shoot architecture in the BTx623 x IS3620C RIL population is *Dwarf3* (*Dw3)*, a sorghum dwarfing gene on chromosome 7 at 59.8 Mbp. The parents of the imaged RIL population, BTx623 and IS3620C, are fixed for non-functional and functional forms, respectively, of the *Dw3* gene which encodes an auxin efflux protein that has pleiotropic effects on stem elongation and additional architecture traits like leaf angle (Multani et al., 2003; Truong et al., 2015). A significant association between *Dw3* and shoot cylinder height is not observed until the second timepoint (34 DAP) while different alleles of *Dw3* introduce significant variability in leaf angle by the earliest timepoint (27 DAP). This is likely because *Dw3* impacts height by impacting stem elongation, and the stem has not yet begun to elongate substantially by the earliest timepoint; as such, the non-functional *dw3* allele caused smaller leaf angles prior to any significant effect on stem elongation (Multani et al., 2003; Truong et al., 2015). Similar to *Dw3*, the effects of *Dwarf2* (*Dw2*), a sorghum dwarfing gene on chromosome 6 near 42 Mbp (but not yet cloned), are significantly associated with shoot cylinder height after the first timepoint (DAP 34, 39, and 44); unlike *Dw3*, *Dw2* is not significantly associated with any other pleiotropic effects on leaf morphology. However, an interval distinct from *Dw2* is observed on chromosome 6 near 51 Mbp for leaf width.

A large interval on chromosome 10 was significantly associated with variability in leaf length and surface area, as well as shoot cylinder height. While the LOD-2 intervals for these traits overlapped when comparing all phenotype by timepoint combinations, the LOD-2 interval for leaf surface area at DAP 39 was distinct from any shoot cylinder height intervals. Additionally, the significant association of the interval with shoot cylinder height is lost after DAP 34, while the association is maintained with leaf traits throughout the timecourse, suggesting that multiple QTL that regulate shoot architecture are present on chromosome 10 (Table S1).

An interval on chromosome 4 was associated with multiple leaf traits, including length, width, and surface area, measured as the average value of leaf numbers 3, 4, and 5 when counting green leaves starting from the bottom of the plant at the time of acquisition. Further analysis showed that plants with BTx623 alleles of an indel marker at the leaf length maximum LOD (MLOD) coordinate (chromosome 4, 62.45 Mbp) had a leaf length of 50.1 cm when averaged across the four timepoints. This was 5.6 cm larger than plants with IS3620C alleles, which had a leaf length of 44.5 cm when averaged across the four timepoints. Additionally, the platform captured changes in leaf length over time; plants with BTx623 alleles increased from an average length of 44.2 cm to an average length of 54.8 cm over the 17 days whereas plants with IS3260C alleles had leaves that increased from an average length of 40.1 cm to an average length of 47.5 cm (Figure 4).

**Figure 5:**
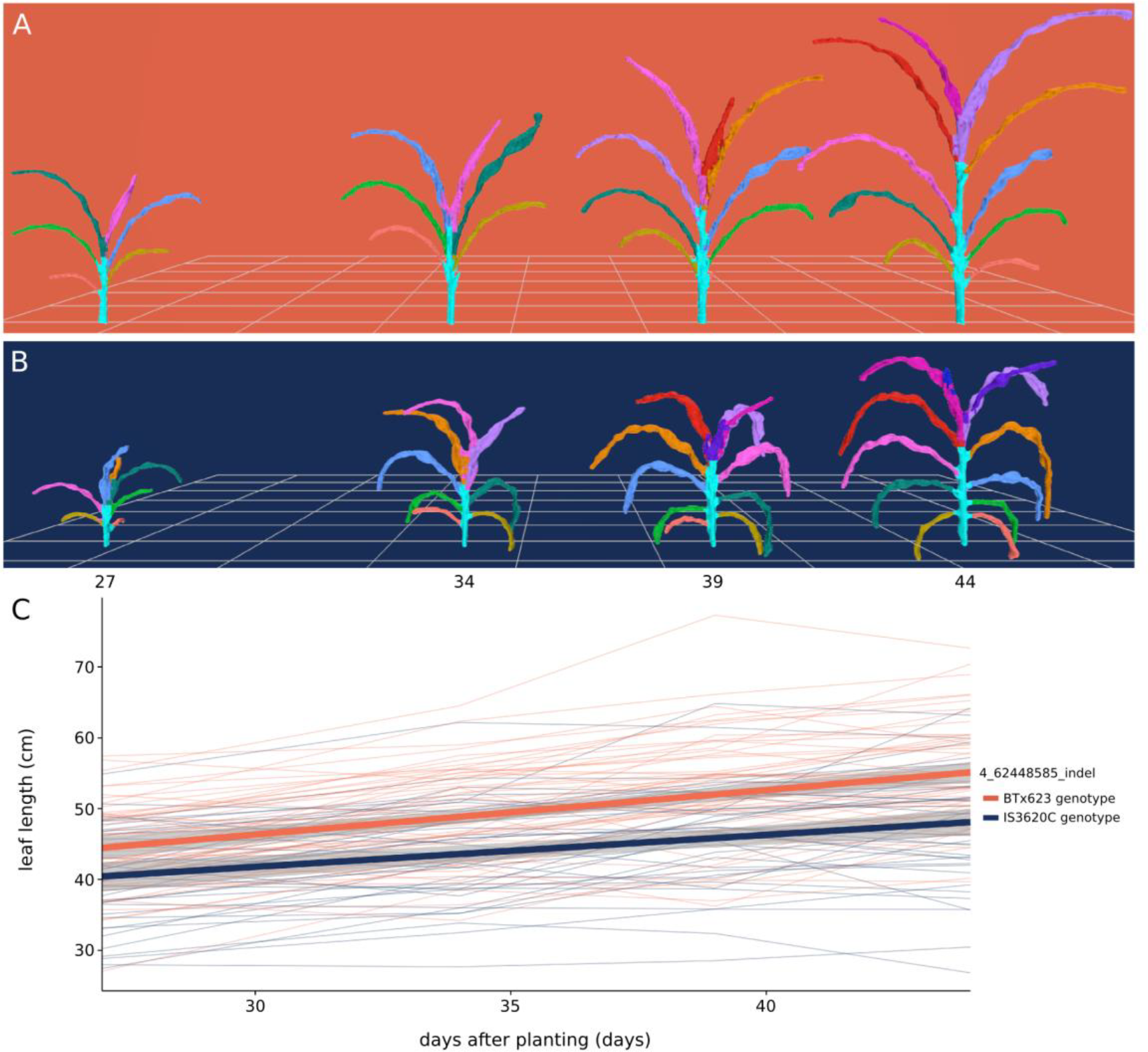
Organ-level measurement of average leaf length over time. (A, B) Meshes displaying development over time for a plant bearing BTx623 alleles (A; RIL 257) and a plant bearing IS3620C alleles (B; RIL 306) of an indel marker on chromosome 4 that had the MLOD for leaf length. (C) Change in average leaf length over time. Each thin line in the plot represents the average leaf length of a RIL (n=3) colored by its genotype. Leaf length was calculated as the average of the third, fourth, and fifth leaves counting from the bottom, corresponding to the light green, dark green, and blue leaves in panels A and B. The two thick lines represent a linear fit for each genotype and 95% confidence intervals.

Because segmentation-dependent traits represent organ-level traits that are often manually measured, QTL identified via the image-based platform for organ-level traits were compared with QTL identified previously for similar traits in the BTx623 x IS3620C population and previous reports on the sorghum dwarfing loci *Dw2* and *Dw3* (Hart et al., 2001; Feltus et al., 2006; Brown et al., 2008; Mace and Jordan, 2011; Morris et al., 2013; Higgins et al., 2014). Most of organ-level QTL intervals found in the present study overlap with comparable or related traits from previous field studies (Table 2). Of note, some of the intervals, like chromosome 6 near 51 Mbp and chromosome 4 near 62 Mbp, may have multiple genes that each affect different traits or genes with pleiotropic effects since these intervals were associated with diverse leaf morphology traits across the studies. Additionally, the genes involved could be environmentally responsive since related but different traits were associated for the intervals when comparing the greenhouse-based and field-based studies (e.g., leaf length in the present study vs. leaf pitch, but not leaf length, in the previous study where leaf pitch measures the length of the leaf from the leaf base to the apex of the naturally-curved leaf). Overall, there was extensive overlap between the QTL intervals identified in previous work and those identified using the imaging platform, suggesting that these genomic loci exert phenotypic effects across multiple studies and environments.

**Table 2:** Comparison of QTL intervals identified using image-based phenotyping with previously reported QTL intervals in the literature. Most QTL intervals identified with the platform overlapped with QTL or causal genes previously previously reported for related phenotypes (Hart et al., 2001; Feltus et al., 2006; Mace and Jordan, 2011; Morris et al., 2013; Higgins et al., 2014). *Dw3* has been previously cloned (Multani et al., 2003). For image-based QTL intervals, the encompassing physical coordinates of the indicated phenotypes across all timepoints were retained as the beginning and end of the interval. The LOD-2 interval and peak coordinate for the phenotype with the maximum MLOD is reported, and the phenotype name is indicated by *. Table S1 contains all identified organ-level QTL. Leaf pitch and leaf curve are both measures of Euclidean distance from the leaf base to the apex of the curved leaf blade, and the leaf base to the leaf tip, respectively (Feltus et al., 2006).

| Chr | Origin | Interval begin (Mbp) | Max MLOD coordinate (Mbp) | Interval end (Mbp) | Related traits with overlapping intervals | Prior locus names |
| --- | --- | --- | --- | --- | --- | --- |
| 4 | Image-based | 57.48 | 62.45 | 63.40 | leaf length*, leaf surface area, leaf width | *QLcv.txs-D2*;<br>*QLpt.txs-D* |
|  | Literature | 61.86 | - | 65.07 | leaf curve, leaf pitch |  |
| 6 | Image-based | 40.10 | 42.77 | 44.83 | shoot cylinder height* | *Dw2* |
|  | Literature | 39.72 | - | 42.64 | pre-flag leaf height |  |
|  | Image-based | 48.45 | 50.97 | 55.08 | leaf width* | *QLcv.txs-I* |
|  | Literature | 53.73 | - | 56.52 | leaf curve |  |
| 7 | Image-based | 59.51 | 59.65 | 59.99 | leaf angle*, shoot cylinder height | *Dw3* |
|  | Literature | 59.821905 | - | 59.829910 | leaf angle, culm height, culm uniformity |  |
| 10 | Image-based | 1.23 | 2.00 | 8.21 | leaf length*, leaf surface area | *QLcv.txs-G*;<br>*QLpt.txs-G*;<br>*QLln.txs-G*;<br>*QHtu.txs.G*;<br>*QHGT_meta1.10* |
|  |  | 5.27 | 7.46 | 52.24 | shoot cylinder height* |  |
|  | Literature | 1.11 | - | 5.76 | leaf curve, leaf pitch |  |
|  |  | 6.40 | - | 13.05 | leaf length, culm height, culm uniformity |  |

In addition to capturing components of plant architecture like leaf morphology, the image-based measurements also capture overall architecture traits that integrate component traits. These composite measurements are difficult or impossible to measure by hand and integrate how component traits interact to influence overall plant architecture and ultimately how a plant canopy intercepts solar radiation. One specific example of such a measurement is shoot compactness, measured as the surface area of the convex hull ofa plant mesh. Shoot compactness is influenced by factors like the leaf angle, the height and planarity of a plant (Figure S4). Accordingly, a strong association between *Dw3* and shoot compactness is present at all timepoints due to the consistent effects of *Dw3* on leaf angle and later effects of *Dw3* on stem growth (Figure 5). As such, composite traits represent measures of overall plant architecture and integrate the interrelationships between component phenotypes. Additional composite traits examined were shoot surface area, shoot center of mass, and shoot height as described in Table 1.

**Figure 6:**
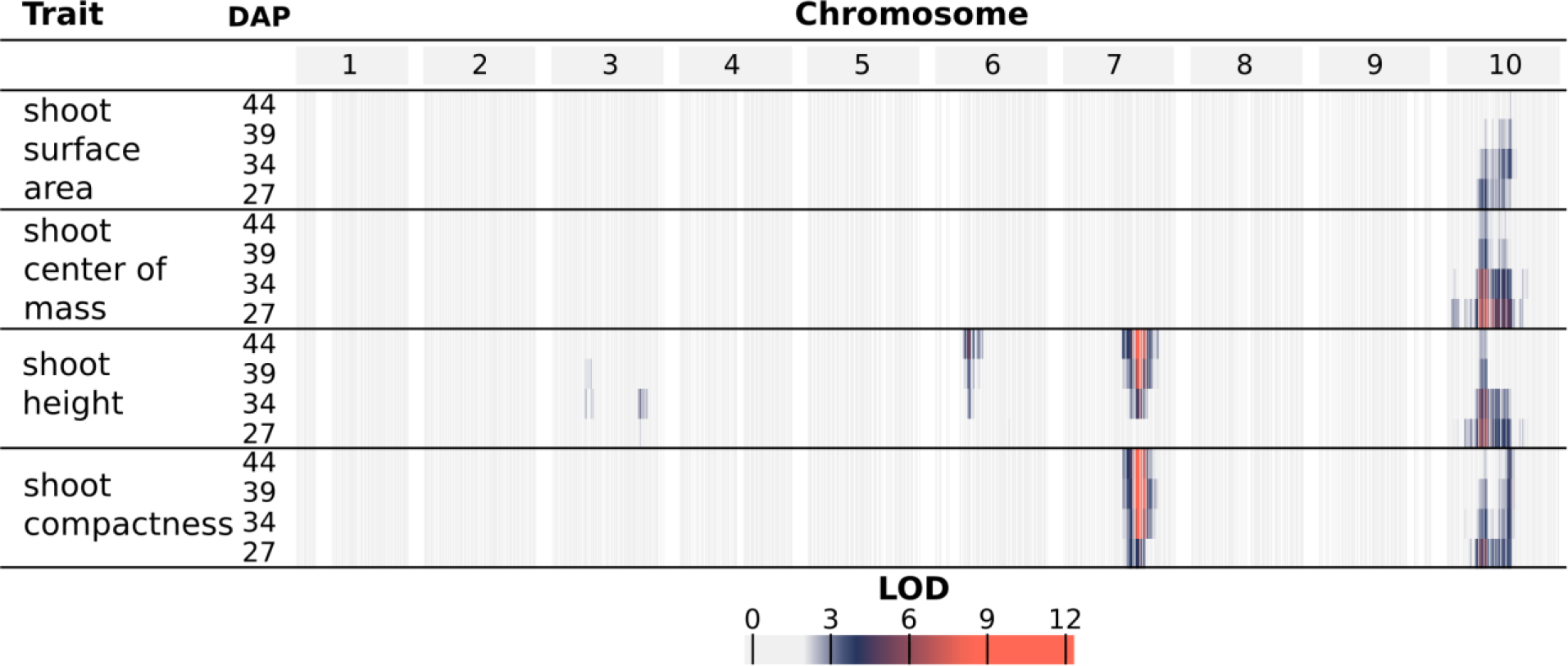
LOD profiles for composite traits. For each phenotype, LOD profiles are based on chromosome-wide scans of chromosomes with QTL based on the most likely multiple-QTL models found by model selection (Figure S7). Each row represents a different trait, and within each trait are four nested rows that each represents a different timepoint (days after planting; DAP). Each group of columns represents a chromosome, and each column represents a marker at its genetic position. Cells are colored by marker LOD for the phenotype at the particular timepoint.

QTL mapping of the selected composite traits identified four genomic intervals with significant contributions to phenotypic variability (Figure 6, Figure S7, and Table S2). Since composite traits are expected to be driven by phenotypic variation in their component traits (and thus correlated) the composite trait QTL are discussed in the context of organ-level QTL with shared intervals. All of the composite traits were significantly associated with a large interval on chromosome 10 at early stages of development (DAP 27 and 35). Consistent with the observation of non-overlapping QTL intervals for organ-level traits of leaf morphology and shoot cylinder height on chromosome 10, at least two QTL are likely present in the interval; canopy compactness is a trait influenced by both leaf morphology and shoot height, and distinct LOD peaks were observed, one at 6 Mbp and one at 52 Mbp (Table S2).

Interestingly, one interval unique to the composite trait measurements was identified on chromosome 3 near 66 Mbp for shoot height, indicating that there are additional component traits driving variability in overall architecture that remain to be resolved and explained by organ-level traits. Alternatively, the impact of the QTL on individual, organ-level traits is relatively small, and only the combined effect across multiple individual traits provide sufficient power for detection. As such, these composite traits represent a useful approach for detecting novel genetic loci.

Due to the effect of *Dw3* on shoot cylinder height and leaf angle, a strong association is present for plant height, and shoot compactness at the *Dw3* locus; likewise, *Dw2* is associated with shoot height. To further quantify the influence of *Dw3*, the shoot height of individuals bearing different alleles of an indel marker near *Dw3* were compared. Plants that have the dominant, functional *Dw3* allele increase in height from, on average, 60.2 cm to 112.6 cm over the 17 day imaging interval, and plants with non-functional *dw3* alleles increase in average height from 56.8 cm to 93.2 cm (Figure 7). Fitting the data to a linear model, *Dw3* plants grew vertically at a rate of 3.1 cm per day whereas *dw3* plants grew at a rate of 2.2 cm per day between 27 days and 44 days after planting. Non-destructive, image-based phenotyping combined with high-throughput genotyping has great potential for parametrizing plant functional-structural modeling and performance prediction with genotype-specific rates of growth.

**Figure 7:**
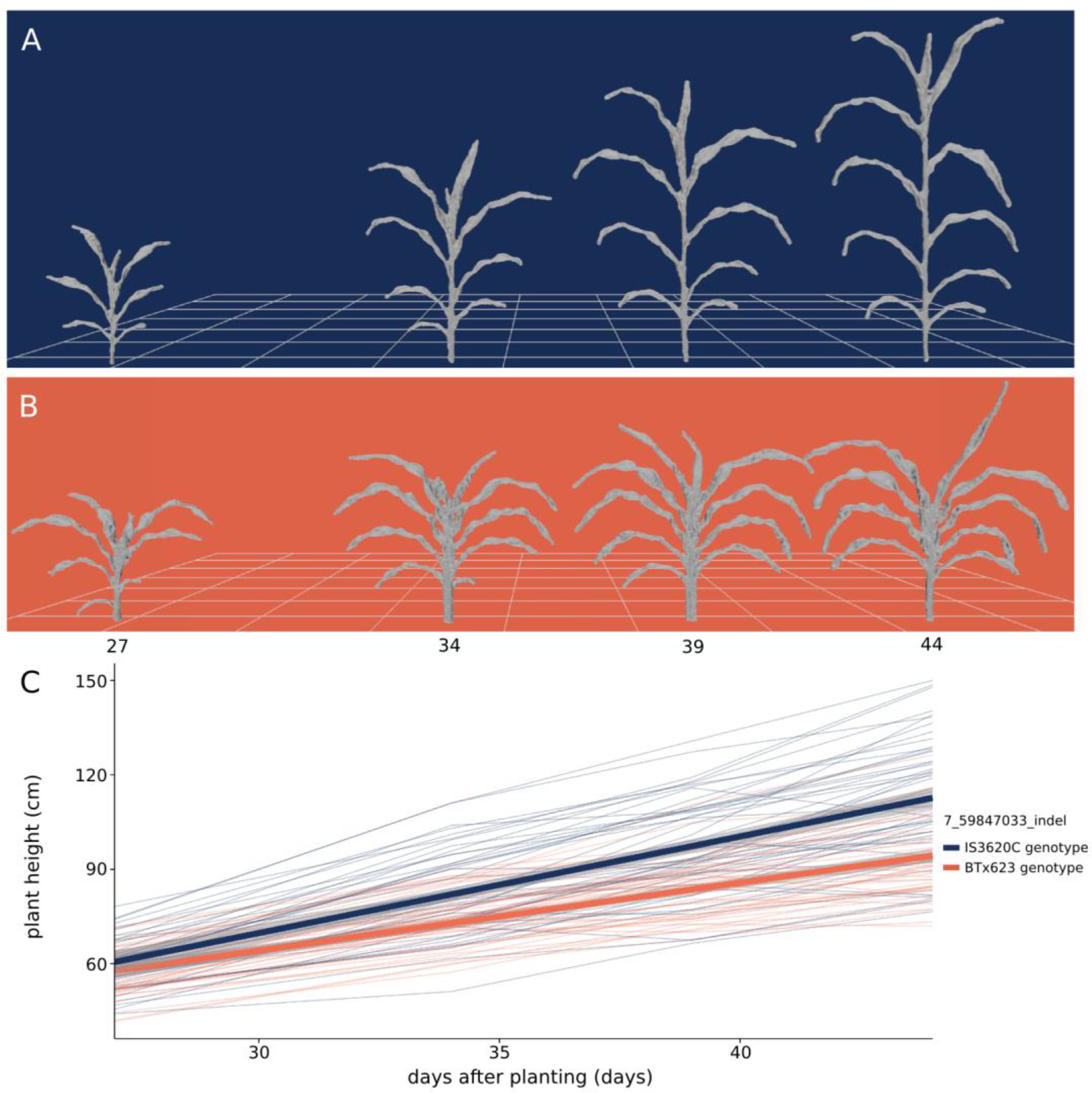
Composite measurement of shoot height over time. (A, B) Meshes displaying development over time for a plant bearing IS3620C alleles (A; RIL 175) and BTx623 alleles (B; RIL 19) of an indel marker closely linked with the *Dw3* gene, an auxin transporter that regulates plant height. (C) Change in plant height over time. Each thin line in the plot represents the average height of a RIL (n=3) colored by its genotype at the *Dw3* locus. Shoot height was measured as the vertical distance from the lowest shoot point to the highest shoot point, including leaves and the inflorescence (Table 1). The two thick lines represent a linear fit for each genotype at *Dw3* and 95% confidence intervals.

## Discussion

A time-of-flight depth camera was used to image sorghum plants from a RIL population, and we developed an image processing pipeline to reconstruct 3D sorghum plants and make automated measurements from the reconstructions. Measurements made in this manner are sufficiently rapid and accurate to enable the identification of multiple genetic loci regulating shoot architecture. As such, we demonstrate that depth imaging represents a useful approach for high-throughput phenotyping of crop plant architecture for the genetic dissection of complex traits.

While the platform successfully identified QTL regulating sorghum architecture (Figures 4 and 6), a number of improvements will be necessary prior to its applicability in even larger scale studies. First, the acquisition process will need to be improved. Registration artifacts were a recurring problem introduced by non-rigid transformations of plant leaves caused by leaf shaking on the turntable, the registration methods used, and sensor noise in acquisition. Multiple potential solutions for these are available, including the use of a registration algorithm capable of handling non-rigid transformations (Zheng et al., 2010; Bucksch and Khoshelham, 2013; Brophy et al., 2015), the use of multiple sensors, the use of real time mesh construction procedures like the Kinect Fusion to average sensor data and rapidly reconstruct the plant (Izadi et al., 2011), or the use of a model-based approach to fit a geometric plant model to the acquired points (Quan et al., 2006; Ma et al., 2008). Second, the segmentation procedure will need to be improved to better distinguish leaves that are in contact with one another, to better automatically identify the shoot cylinder of the plant, and to potentially make it applicable to other grass or plant species. General algorithms that can accurately segment plant organs for images or meshes from multiple species will be of value and will need to scale to complex field scenes involving multiple plants and heavy occlusion. Mapping additional data types, such as visible, infrared, or hyperspectral, onto the point clouds will also be of value for both controlled-environment and field applications.

A major benefit of image-based phenotyping is its non-destructive nature because insight into the temporal onset of genetic regulation is valuable in dissecting its mechanistic basis. Markers tightly linked with *Dw3*, a gene encoding an auxin transporter, are associated with leaf inclination angle, plant compactness, and canopy length prior to their association with shoot height and shoot cylinder height, suggesting that changes in auxin transport caused by different *Dw3* alleles are introducing variability in leaf development and overall shoot compactness prior to large effects on stem elongation (Figures 4, 6, and 7). Additionally, variation in the shoot cylinder height at the earliest timepoint is most associated with an interval on chromosome 10 (Figure 4). This QTL is the primary driver of variability in shoot height and shoot cylinder height until the variability in stem growth introduced by alleles of *Dw2* and *Dw3* increases, and it may be related to the timing of a developmental transition (Figures 4 and 6). It is likely that multiple QTL are present on chromosome 10 given that distinct LOD peaks at 2 Mbp, 7 Mbp, and 52 Mbp were observed; additional experimentation will be necessary to resolve the contributions and temporal prevalence of specific QTL in the interval.

Many of the QTL identified via image-based phenotyping overlapped with QTL for comparable traits discovered in prior field experiments (Table 2). These shared QTL represent good candidates for continued investigation as they display robust phenotypic effects across multiple experiments and conditions. Notably, despite sharing overlapping intervals, the associated traits sometimes differed. For example, the present study identified significant associations between leaf length, width, and surface area with an interval on chromosome 4; a similar interval was identified in previous work for leaf curve and leaf pitch, but was not significantly associated with leaf length in the previous study (Table 2). While all of these traits are aspects of leaf morphology and share relationships, additional experimentation will be necessary to determine whether these represent one QTL with pleiotropic effects (as is observed with *Dw3*), one QTL with different environmental responses, different QTL with overlapping intervals, or some combination of these possibilities.

## Conclusions

Depth imaging and subsequent processing enabled the rapid acquisition of multiple shoot architecture phenotypes from a sorghum QTL mapping population, and genetic loci contributing to variation in shoot architecture were identified. Depth cameras represent a practical tool for rapidly measuring plant morphology, and their applications to plant phenotyping alongside other imaging modalities will be useful for both controlled-environment and field phenotyping scenarios. Integrated platforms that merge image-based phenotyping approaches, genetics, and performance modeling will enable rapid improvements in understanding plant biology and will promote the selection and engineering of plants for superior performance in target applications.

## Acknowledgements

The authors thank Sergio Hernandez for his assistance during image acquisition. We thank Marcin Kalicinski for RapidXML which is used to parse configuration files during mesh processing (http://rapidxml.sourceforge.net/; accessed February 2016). We thank the Blender Foundation for the 3D modelling and rendering package, Blender, which was used to stage meshes for the figures (https://www.blender.org/; accessed February 2016). We thank the Texas A&M Institute for Genome Sciences and Society for maintaining the TIGSS HPC Cluster which was used to calculate main effect, heavy chain, and light chain penalties for multiple-QTL mapping. Finally, we thank the anonymous reviewers for providing constructive feedback during the review process.

